# The impact of incongruence and exogenous gene fragments on estimates of the eukaryote root

**DOI:** 10.1101/2021.04.08.438903

**Authors:** Caesar Al Jewari, Sandra L. Baldauf

**Affiliations:** Program in Systematic Biology, Department of Organismal Biology, Uppsala University, Uppsala, Sweden 75236

**Keywords:** phylogenomics, congruence, eToL, eukaryote root, Discoba, mitochondrial proteins

## Abstract

Phylogenomics uses multiple genetic loci to reconstruct evolutionary trees, under the stipulation that all combined loci share a common phylogenetic history, *i*.*e*., they are congruent. Congruence is primarily evaluated via single-gene trees, but these trees invariably lack sufficient signal to resolve deep nodes making it difficult to assess congruence at these levels. Two methods were developed to systematically assess congruence in multi-locus data. Protocol 1 uses gene jackknifing to measure deviation from a central mean to identify taxon-specific incongruencies in the form of persistent outliers. Protocol_2 assesses congruence at the sub-gene level using a sliding window. Both protocols were tested on a controversial data set of 76 mitochondrial proteins previously used in various combinations to assess the eukaryote root. Protocol_1 showed a concentration of outliers in under-sampled taxa, including the pivotal taxon Discoba. Further analysis of Discoba using Protocol_2 detected a surprising number of apparently exogenous gene fragments, some of which overlap with Protocol_1 outliers and others that do not. Phylogenetic analyses of the full data using the static LG-gamma evolutionary model support a neozoan-excavate root for eukaryotes (Discoba sister), which rises to 99-100% bootstrap support with data masked according to either Protocol_1 or Protocol_2. In contrast, site-heterogeneous (mixture) models perform inconsistently with these data, yielding all three possible roots depending on presence/absence/type of masking and/or extent of missing data. The neozoan-excavate root places Amorphea (including animals and fungi) and Diaphoretickes (including plants) as more closely related to each other than either is to Discoba (Jakobida, Heterolobosea, and Euglenozoa), regardless of the presence/absence of additional taxa.

## Introduction

Over the past twenty years, large multigene (phylogenomic) datasets have confidently and consistently identified the broad outlines of the eukaryote tree of life (eToL) (Baldauf et al. 2000; Bapteste et al. 2002; Brown et al. 2018; Burki et al. 2016, 2020; Strassert et al. 2019). As a result, nearly all known eukaryotes can be assigned to one of three supergroups: Amorphea, Diaphoretickes and Excavata. Amorphea includes animals (Holozoa), fungi (Holomycota or Nucletmycea), amoebozoan amoebas (Amoebozoa) and several less well characterized lineages. Diaphoretickes encompasses nearly all other well-known eukaryotes including land plants and nearly all of the algae interspersed with numerous non-photosynthetic lineages such as ciliates, oomycetes and apicomplexan parasites. Excavata is by far the least well known and most poorly understood of the supergroups and is thought to consist mainly of Discoba, Metamonada, and, possibly Malawimonadida (Adl et al. 2018). Within Excavata only the monophyly of Discoba (Jakobida, Euglenozoa and Heterolobosea) is well-established (Kamikawa et al. 2014). Discobans are largely free-living and have functional, if molecularly diverse mitochondria, while all known metamonads are micro- or anaerobic and lack mitochondrial DNA (mtDNA) and nearly all mitochondrial proteins (Hjort et al. 2010).

One of the most important outstanding questions in eukaryote phylogeny is the position of the root. Since eukaryote genomes are mosaics with roughly half of their genes of clearly archaeal or bacterial origin (Brueckner & Martin 2020; Cotton & McInerney 2010), eToL can be rooted with either bacterial or archaeal homologs. Of these, archaeal genes trace to the eukaryote first common ancestor (FECA), while bacterial genes trace to the origin of endosymbiotic organelles. Since mitochondria arose after FECA but before the eukaryote last common ancestor (LECA; Gabaldón 2018), this makes bacteria a potentially closer outgroup to eukaryotes than archaea. Nonetheless, rooting eToL with bacteria is complicated by the facts that only ~10% of mitochondrial proteins (mitoPs) are shared across eukaryotes (Gray 2012), and these proteins trace to different groups of bacteria, although mostly alpha- and gamma-proteobacteria (αP_bac and γP_bac, respectively) (Kurland & Andersson 2000). In addition, Discoba are the only excavates with mitoPs meaning that an eToL rooted with bacteria can include only one representative of the diverse and not necessarily monophyletic Excavata (Karnkowska et al. 2016). Thus, an eToL based on mitoPs is only one step in defining the eukaryote root, albeit an important one.

Previous attempts to recover the eToL root with mitoP data are mainly the work of Derelle and Lang 2012 (DL2012), He et al. 2014 (H2014) and Derelle et al. 2015 (D2015). These studies used partially overlapping subsets of mitoPs and similar taxa but reached different conclusions. DL2012 and D2015 recovered the more traditional unikont-bikont root (Amorphea sister), while H2014 recovered a novel neozoan-excavate root (Discoba sister). These studies differed mainly in outgroup selection (αP_bac only - DL2012/D2015, versus mainly αP - and γP_bac - H2014) and inclusion (DL2012/D2015) or exclusion (H2014) of Malawimonads. In addition, the two studies differ in phylogenetic method, with DL2012 and D2015 using site-heterogeneous evolutionary models and removal of fast evolving site (FSR), while H2014 used site-homogeneous models with no FSR and a site heterogeneous model with a covarion process. Both studies also reported evidence of substantial incongruence in the other data sets.

Congruence is a core premise of phylogenomics, *i*.*e*., the assumption that the loci to be combined for phylogenetic analysis share the same underlying evolutionary history (Huelsenbeck et al. 1996; Roger et al. 2012). If this premise is met, then increasing data should amplify a weak common signal (Kupczok et al. 2010; Philippe et al. 2011; Wägele et al. 2009). Thus, assessing congruence across a multi-gene dataset is critical. This is generally done by evaluating single gene trees (SGTs) by either visual or automated inspection.

However, SGTs generally lack sufficient phylogenetic signal to resolve the deepest branches in a tree, making it particularly difficult to assess incongruence at deep levels. Variation in length and degree of conservation among loci also make it challenging to apply a uniform standard evaluation process for SGTs. Thus, evaluating SGTs individually becomes a laborious and troublesome process for large numbers of loci.

The primary source of molecular incongruence is when genes arising by ancient duplication (out-paralogs) or by horizontal transfer (xenologs) are mistaken for strictly vertically inherited genes (orthologs), since only orthologs directly track organismal phylogeny. Strategies for tackling the congruency problem can be classified into two general categories: 1) detecting persistent outliers by measuring the deviation from a global reference point or 2) clustering genetic loci into subsets based on a compatibility score. Both approaches require a metric to measure the compatibility scores, whether between loci (partitions) or between partitions and a global reference point. This metric is usually based on tree dissimilarity (distance), tree likelihood, or a combination of the two (Smith et al. 2020).

The most widely cited methods for automated testing of congruence are Conclustador (Leigh et al. 2011) and Phylo-MCOA (De Vienne et al. 2012). Conclustador uses a distribution of bootstrap or posterior trees for each locus. It then measures the distance between each pair of loci by counting the number of shared bipartitions in their tree set, after removing non-shared taxa. Phylo-MCOA (De Vienne et al. 2012) uses multiple co-inertia analysis to identify outliers at two levels: complete outliers (entire loci or species) and cell by cell outliers (specific loci for specific species). Missing data are filled in with average values from across the data. The method uses only a single tree for each locus, so the statistical significance of topological differences is not taken into account.

We have developed two algorithmic approaches to assess incongruence in multi-gene data. Protocol_1 is a combinatorial method using gene jackknifing to rank deviation from a central mean. The ranking is measured by quantifying topological inconsistencies in data subsamples and extracting sequence-specific deviation. The problem of missing data is overcome by requiring each jackknife sample to include all taxa. The second method uses a sliding window to assess inconsistent signal at the sub-gene level using a pre-selected set of target taxa to narrow the search space. Both methods are applied to a set of core mitoPs previously used to test the eukaryote root.

## Materials and Methods

### Data sources

Sequences were obtained primarily from GenBank (Benson et al. 2013) and iMicrobe (Youens-Clark et al. 2019). Additional sources were used for Amoebozoa (Kang et al. 2017; provided by the author), Dictyostelia (Fey et al. 2013), and three Discoba (*Acrasis kona*, *Andalucia godoyi*, *Seculamonas ecuadorienses*; Fu et al. 2014; He et al. 2014). Taxa were selected to give broad and balanced taxonomic coverage of the main eukaryotic groups and the main bacterial outgroups, with a preference for organisms with higher sequence coverage. The complete data set includes 208 organisms (Supplementary Table S1).

### Data assembly

Sequences were identified by BLASTp, and all hits below an e-value of 1e-5 were retrieved. The BLAST queries consisted of all 75 proteins used in various combinations in three previous studies (Derelle et al. 2015; Derelle & Lang 2012; He et al. 2014). Homologs for each protein were searched for separately using probes from three different organisms, which together represent the three major divisions of eukaryotes - *Naegleria gruberi* (Excavata), *Capsaspora owczarzaki* (Amorphea) and *Selaginella moellendorffii* (Diaphoretickes). The retrieved protein sequences were aligned using Mafft 7.450, with different algorithms depending on the number of sequences (Katoh & Standley 2013); L-INS-i --maxiterate 1000 was used for alignments of ≥250 sequences and FFT-NS-i --maxiterate 1000 otherwise. The resulting alignments were trimmed using the gappyout option in trimAl 1.4 (Capella-Gutiérrez et al. 2009). Visual inspection identified 21 alignments in which automated trimming either removed too many columns or left too many gaps to be useful for sliding window analysis (Protocol_2, below); these alignments were instead trimmed by hand for Protocol_2 analyses only.

SGT analyses used individual protein alignments, which were iteratively screened for potential horizontal gene transfer (HGT) and out-paralogs by multiple rounds of phylogenetic analysis using FastTree (-gamma-lg-spr 6-mlacc 2-slownni-slow; Price et al. 2010). This was followed by a final more stringent phylogenetic assessment using RAxML-NG with the LG+GAMMA model and standard bootstrapping (Kozlov et al. 2019). The screening resulted in seven proteins being rejected because they either showed strong support for major conflicts with established eukaryote phylogeny (Adl et al. 2018), indicative of HGT or deep paralogy, or lacked representation of one of the three major eukaryote groups. A further eight alignments were found to consist of two strongly separated eukaryote-universal paralogs (100% bootstrap support), and these were each split into two separate alignments. The result is a final dataset of 76 proteins and 180 taxa.

### Congruence tests – Protocol_1

Protocol_1 assesses data outliers through random subsampling of partitions (jackknifing; Fig. 1). The three mains steps are 1) subsample assembly, 2) tree inference, and 3) tree evaluation. Step 1 generates the subsamples and an inclusion table that tracks the partition composition of each subsample (Fig. 1a). The subsamples are created at random without replacement with two requirements: i) each sequence must be present in an equal number of subsamples and ii) each subsample must contain at least one sequence for every taxon. The first requirement is achieved by randomly splitting the dataset into subsets with equal numbers of partitions. This is repeated until reaching the desired number of subsets while satisfying the equal inclusion frequency requirement. The equal inclusion requirement means that the chosen number of subsamples must be divisible by the total number of partitions. The size of subsample, in turn, depends on the taxon-wise ratio of missing data, i.e., the ratio of missing versus available markers for each taxon. If one or more taxa have a high ratio (roughly, >50% missing), it may be necessary to either increase the subsample size or to exclude the highly-incomplete taxon in order to satisfy the second requirement.

**Figure 1.**
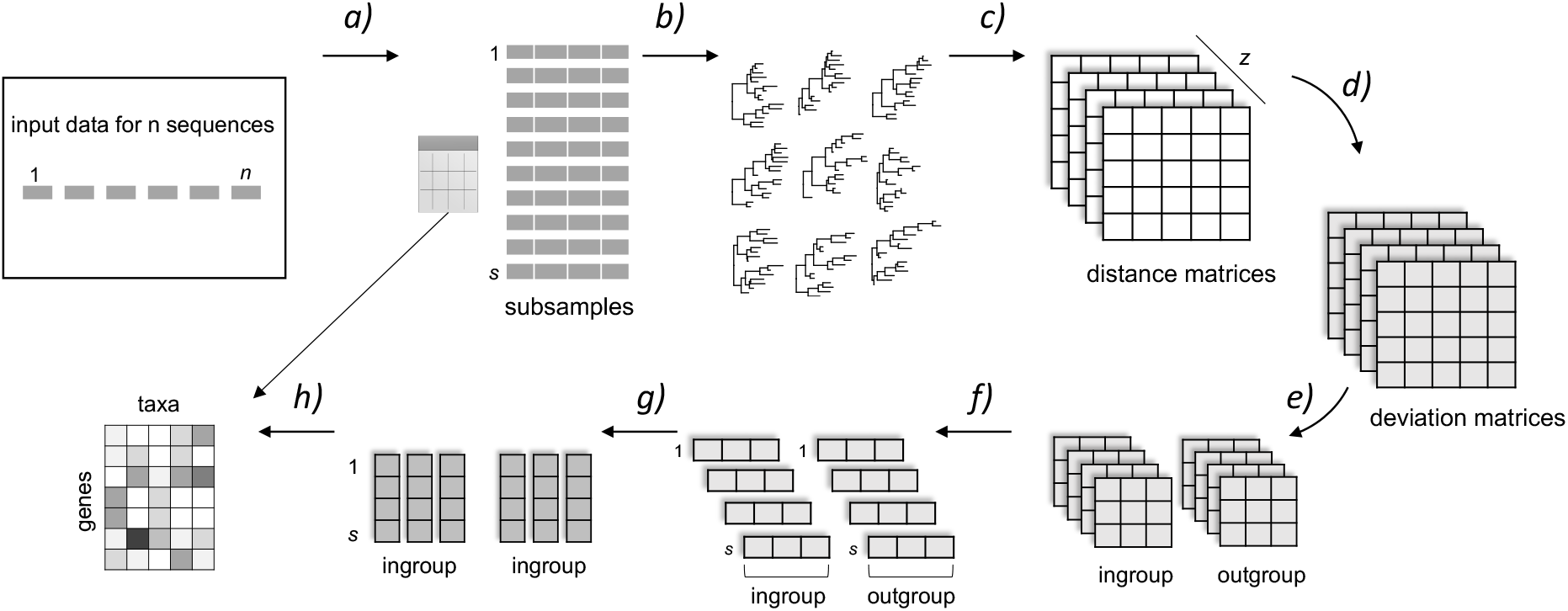
Protocol_1: Estimating phylogenetic incongruence scores in multigene data by random sub-sampling (jackknifing). The protocol consists of 8 main steps. a) Random balanced subsampling of ‘n’ sequences generates ‘s’ subsamples, with sequence distribution across subsamples recorded in an inclusion table. b) Maximum likelihood trees are estimated for each subsample, and c) each tree is converted into a nodal distance matrix. d) Tree deviations are estimated with reference to the inner mean across the ‘z’ axis. e) Matrices are split into ingroup and outgroup to be assessed separately, and f) a sum of deviations is calculated for each taxon by summing across columns. g) Taxon-gene values are scaled by multiplying by the mean of the lowest 10% values across all subsamples. h) The inclusion table from step 1 is used to convert scaled values from step 7 into a single scoring matrix. In an optional final step, the scoring matrix may be converted into a heat map using additional scripts. All scripts are available from https://github.com/caeu/, except for the heatmap script which is available upon request.

Step 2 of Protocol_1, the only computationally demanding step, generates a maximum likelihood (ML) tree for each subsample by ML analysis using RAxML-NG 0.9 and the LG substitution model (Fig. 1b). Unlike steps 1 and 3, which can be executed on a desktop computer, step 2 is better suited for multicore parallel processing. Step 3 uses the output trees from step 2 to calculate sequence-specific outlier scores. The script then masks sequences with the top 10% highest scores, and outputs the resulting masked (trimmed) concatenated dataset (Fig 1c-h). Outlier scores are calculated by first converting each tree to a pair-wise nodal distance matrix (Fig 1c). Deviation values are then calculated for each taxon-pair for each distance matrix from the calculated truncated mean at 35% for all taxon-pair distances across all matrices according to equation 1:

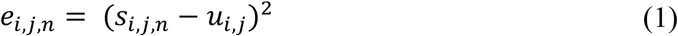

where *s_i,j,n_* is the distance between the taxon pair (i,j) at the *n*th subsample, and *u_i,j_* is the truncated mean for the taxon pair (i,j) for the subsamples (Fig. 1d). Deviation scores are then rescaled within each matrix by squaring each deviation value and dividing it by the mean of all deviation values in that matrix. This gives higher weight to deviations in trees with fewer but larger errors.

The deviation matrices are then split into ingroup and outgroup submatrices and analysed separately (Fig. 1e). This is a precautionary measure, otherwise optional, to avoid interference from the high phylogenetic noise often present in or arising from the outgroup, which may overwhelm a weaker ingroup signal. The outgroup matrix also includes a single ingroup taxon, specifically the one with the lowest standard deviation as calculated from the ingroup submatrix. This taxon serves as a stable reference point for calculating instability scores for the outgroup taxa.

Finally, cumulative taxon-specific deviation scores are calculated for each subsample by summing the values in their respective deviation matrix (column wise or row wise). The sum of scores of each sequence-taxon data point are then rescaled by multiplying them by the mean of the lowest 10% points across all the data points for that sequence-taxon (Fig. 1g). The inclusion table from step 1 (Fig. 1a) is then used to transfer the sum of deviation scores from the subsamples to the corresponding proteins. Finally, the original data set is modified by masking the sequences that rank in the top 10% deviation scores, followed by masking any taxon or partition with more than 50% missing data and outputting the final filtered dataset. To help visualize the results, the full set of deviation scores across the data set can be converted into a heatmap (Fig. 1h, script available on request).

### Congruence tests – Protocol_2

Protocol_2 uses a sliding window to evaluate orthology at the sub-gene/protein level. The method requires designating a monophyletic group of interest (test taxon) and a phylogenetically conservative and well-resolved taxon set (core taxa) to provide a stable background against which to evaluate the test group data (Fig. 2). The stable group and test group are defined *a priori*, for example based on results from Protocol_1. Data are prepared as above, by alignment and trimming of individual orthologous sequence sets. Alignments are then inspected visually to identify windows missing all data from either the test or core taxon groups, and these windows are then deleted for Protocol_2.

**Figure 2.**
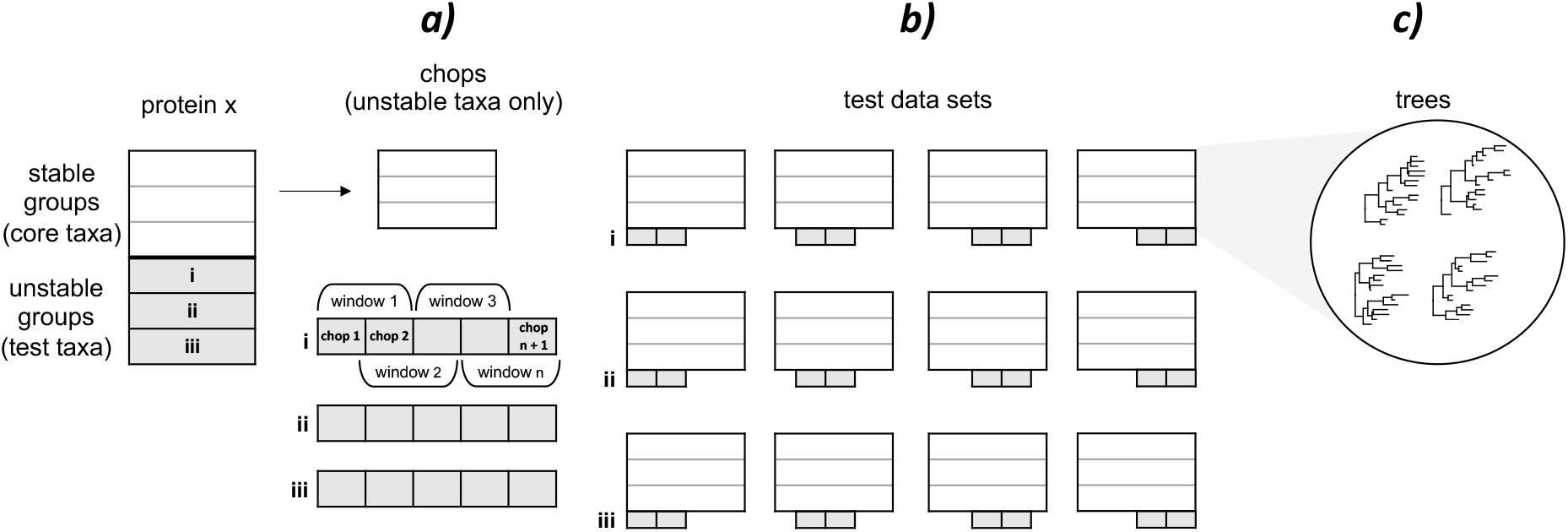
Protocol_2: Sliding window assessment of congruence for protein fragments. The example shown is for testing fragment incongruence in three taxa (“test taxa”), which can be single taxa or clades of related taxa. A taxonomically broad set of stable taxa (“core taxa”, selected *e.g*, based on Protocol_1) provide a stable background against which to test congruence. a) Each protein is broken into 2-5 equal size fragments or “chops”, depending on protein size according to equation 1. Adjacent chops are then combined pairwise to create semi-overlapping windows (50% overlap). In this example, the protein is divided into 5 chops which combine to give 4 overlapping windows. b) Alignments for the core taxa are assembled separately for each window for each of the three test taxa (labelled i-iii) resulting in 12 alignments for a 4-window protein. c) Phylogenetic trees are constructed for each window for each test taxon set using four method-model combinations: RAxML LG, RAxML LG+gamma, IQTREE LG, IQTREE LG+gamma. A taxon-specific window is flagged if its sequence occurs nested within any and the same core taxon clade in all four trees. Chops that are flagged for both underlying windows in which it occurs are scored as potentially incongruent chops (PICs). Scripts are available from https://github.com/caeu/.

The protocol has three main steps: 1) sliding window data assembly, 2) tree inference, and 3) tree evaluation (Fig. 2). Similar to protocol 1, the pipeline implementation assumes running steps one and three on a local computer, while step two requires a multicore system. Step one first divides each trimmed alignment into 3-10 equal size fragments (chops), with the size and number of chops based on alignment length according to the equation:

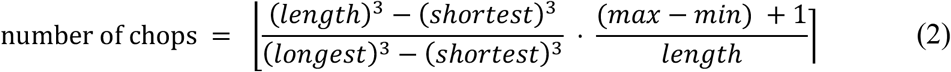

where *length* is alignment length, and *shortest* and *longest* are the shortest and longest alignments in the data set (chop size may vary +/-1 for alignments not precisely divisible by the number of chops). The chops are then combined sequentially in pairs to generate a sliding window across the alignment, with 50% overlap between each chop pair (Fig. 2a). Any window with less than 20 gap-free columns for any major subdivision of the test group is then omitted as these are unlikely to give useful information. A map of the chops’ structure is generated and saved to record which chops are in which data set and to keep track of each set of sliding windows.

Chop pairs are analyzed separately for each division of test taxa by phylogenetic analysis in combination with full alignments for the core taxa (Fig. 2b). Four ML trees are constructed for each sliding window using four different method-model combinations: RAxML-NG and IQ-TREE with LG and LG+Gamma models (Fig. 2c). Chops for which the test taxa are nested within one and the same division of the stable group in all four trees and in two contiguous windows are marked as violating orthology (potentially incongruent chops or “PICs”). Chops at the ends of an alignment or neighboring a missing data window are evaluated based only on the single window in which they occur. In a final step, all PICs are masked from the full data set, and the remaining data are concatenated to produce the final Protocol_2 output. Alternatively, Protocol_2 can be limited to alignments with a minimum of one sequence for each of the test subgroups. This restriction can be applied before and/or after implementing Protocol_2 analysis.

All calculations for both protocols are implemented in R v3.6.3 (R Core Team 2020). The import and transformation of trees and sequences use the Ape package v5.0 (Paradis & Schliep 2019). The R package Rfast is used to speed up parts of the calculations (Tsagris & Papadakis 2018). All scripts are available from https://github.com/caeu/eukRoot.

### Multiprotein Phylogeny

The final mitoP supermatrix was analysed with five different method-model combinations: 1) RAxML NG with the LG+Gamma model and standard bootstraps (Kozlov et al. 2019), 2) IQ-TREE with the LG+Gamma model and ultra-fast-bootstraps (UFB) corrected with the “bnni” option (Hoang et al. 2018), 3,4) IQ-TREE with the C60+LG+G+F model and PMSF bootstraps and also with UFB+bnni (Wang et al. 2018), and 5) PhyloBayes-MP 1.8 (Lartillot et al. 2013). For IQ-TREE C60 analyses, the ML tree and the UFB bootstraps were inferred first using the model, and then the ML tree was used to generate the site-specific frequency profile for the PMSF bootstraps. PhyloBayes was run with two independent chains for 10,000 generations, and a consensus tree and posterior probabilities were calculated from both chains once they had converged.

All analyses other than tree reconstruction were implemented as R Pipeline scripts and executed on a local computer. Tree reconstruction and bootstrapping were run with the computational resources of Uppsala Multidisciplinary Center for Advanced Computational Science (UPPMAX).

## Results

### Data filtration Protocol_1

Two complementary methods have been developed to screen phylogenomic data for potentially problematic components. The first method screens for incongruent taxon-specific loci by identifying and ranking high influence outliers using a whole gene/protein resampling approach (Protocol_1; Fig. 1). The second method uses a sliding window protocol to screen for potentially incongruent sequence fragments (Protocol_2). The two protocols were applied to a data set of 76 mitochondrial proteins (mitoPs) used in various combinations in three previous studies examining the position of the eukaryote root, abbreviated here as D2015, DL2012, and H2014 (Derelle et al. 2015; Derelle & Lang 2012; He et al. 2014).

For Protocol_1 (Fig. 1), initial trials with the 76 mitoP proteins indicated that 14 was the minimum number sufficient to avoid creating subsets with complete missing data for any taxon, while being large enough to produce a stable distribution of well-resolved trees against which to detect aberrations. Each protein was then sampled 364 times, creating a total of 1976 subsamples, which is the closest number to the 2000 sample target that is divisible by 76 (total number of proteins). The resulting phylogenies were used to produce dataset-wide influential outlier scores, displayed here as a heatmap (Fig. 3), after which the sequences in the top 10% of outlier scores were then masked. Since masking increases missing data, the revised data set was trimmed column wise and row wise for excess missing data using a 50% cutoff (Fig. 4). *i.e*., any protein or taxon with greater than 50% missing entries was removed (Fig. 4, black marks along x and y axes). Altogether Protocol_1 reduces the mitoP data from 180 taxa and 76 proteins to 156 taxa and 66 proteins in preparation for phylogenetic analysis (below).

**Figure 3.**
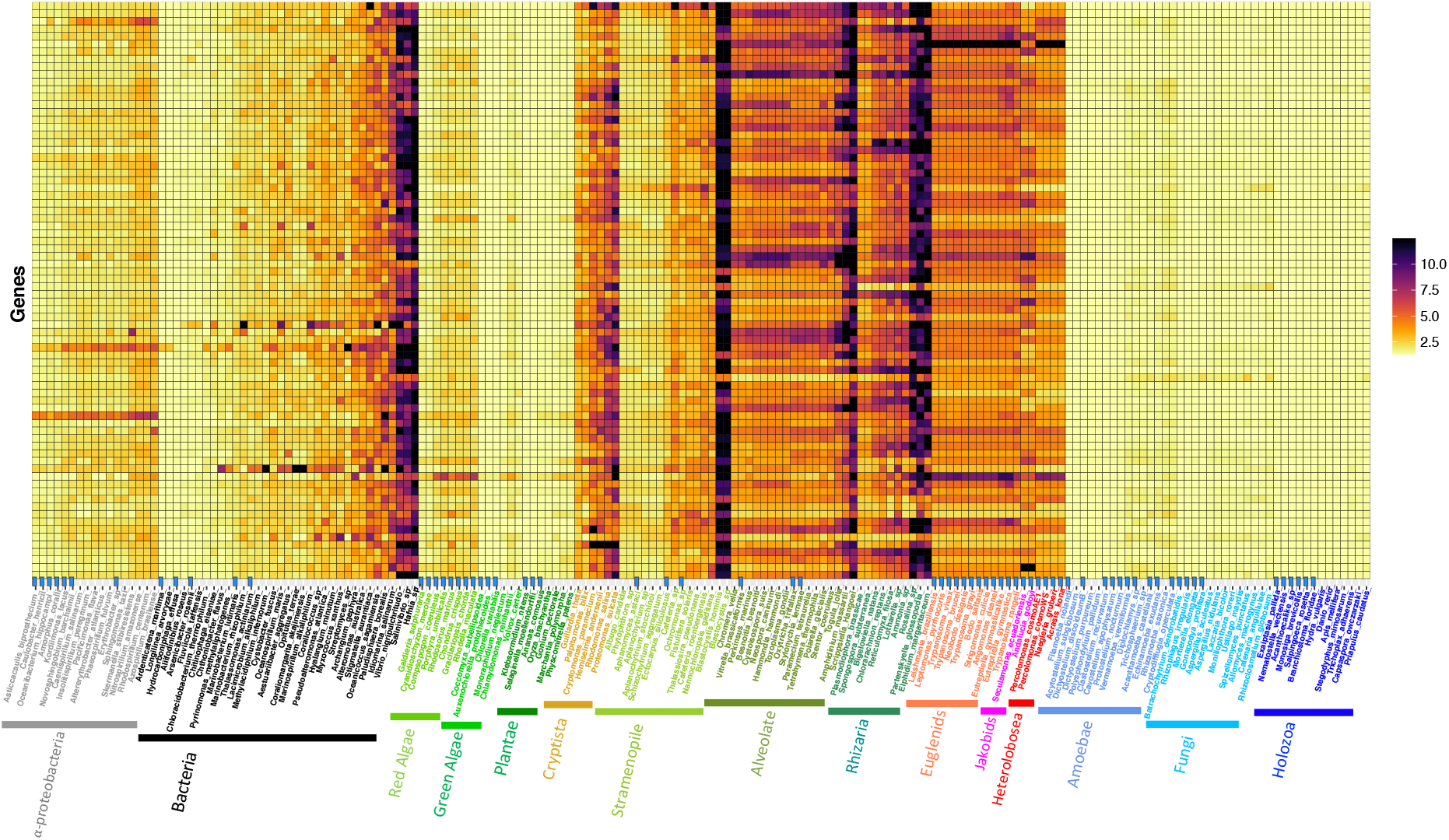
Protocol_1 derived deviation scores for 76 mitochondrial proteins from diverse eukaryotes. The deviation scores for 76 proteins (y-axis) for 180 taxa (x-axis) were determined using the persistent outlier detection method described in Figure 1 (Protocol_1). Deviation score values are indicated by cell color intensity, with darker colors indicating higher deviation according to the scale on the right. Blue flags along the x-axis indicate taxa used in Protocol_2 analyses. Species names are color coded to indicate major taxonomic groups (kingdoms / superkingdoms / domains).

**Figure 4.**
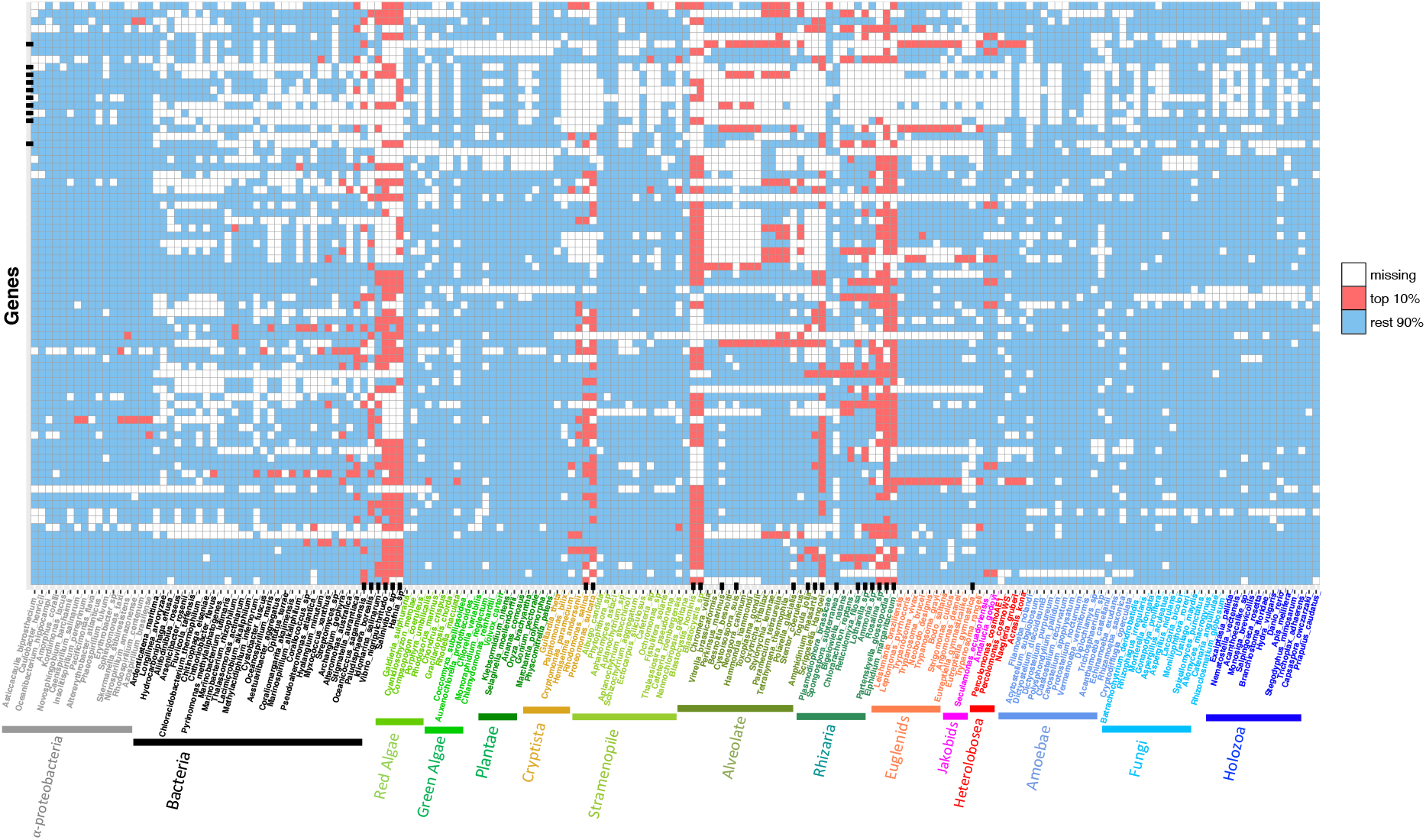
Distribution of missing data and high deviation scores across all mitoP proteins and taxa following Protocol_1 analysis. Presence/absence of sequence data is indicated for 76 mitoP proteins (y-axis) for 180 taxa (x-axis) after Protocol_1 (outlier detection) masking. Cell colors indicate protein presence (blue), absence (white), and whether or not the sequence is among the top 10% ranked outliers identified by Protocol_1 (red). Black flags along the y- and x-axis indicate proteins and taxa, respectively, with more than 50% missing data. Species names are color coded to indicate major taxonomic groups (kingdoms / superkingdoms / domains).

Inspection of the Protocol_1 deviation heatmap shows strong and consistent phylogenetic signal across Amorphea, Plants (Archaeplastida) and most Bacteria for most or all of the mito proteins (Fig. 3). Most of the Stramenopiles are also highly consistent, with the exception of the two *Blastocystis* species, which reflects the fact that these have non-respiring mitochondria (Stechmann et al. 2008). In contrast, strong outliers are seen for many proteins in Alevolata, Rhizaria, Cryptista and Discoba. This is particularly striking for *Amphidinium massaratii* and *Scrippsiella hangoel* (Dinoflagelata) in Alveolata, *Rosalina sp*. and *Elphidium margaritaceum* (Foraminifera) and Partenskyella glossopodia (Cercozoa) in the Rhizaria, and most of the Crpytista. As a result, two of the three major divisions of Diaphoretickes, and most, if not all, of the Cryptista have substantial outlier content distributed widely if unevenly throughout these data (Fig. 3). Most importantly here, the supergroup Discoba shows consistently high deviation scores compared to the rest of the data, with over 60% of Discoba sequences showing deviation scores ranking in the upper quartile. Thus, while Amorphea is nearly completely outlier-free and even Diaphoretickes has largely outlier-free major divisions, particularly Stramenopiles and Plants (red algae, green algae and land plants), no major division of Discoba is largely free of outliers (Fig. 3).

While the excess of outliers in Discoba may be due to some unusual characteristic of the taxon, Discoba is also distinguished as being is by far the least well sampled of the three major divisions of eukaryotes examined here. Thus, it is likely that the consistency seen here in groups such as animals, plants and fungi reflects the fact that they are molecularly well-sampled and well-studied. This makes it easier to identify and avoid outlier taxa while still maintaining the broad taxonomic representation needed to recover deep phylogenetic branches. In fact, there is a strong correlation between breadth of taxonomic sampling in GenBank and Protocol_1 deviation scores (Fig. S1). Unfortunately, avoiding outlier taxa is not possible at present for Discoba, where all currently available genomic data must be used to achieve any semblance of taxonomic breadth. This includes using molecularly highly divergent taxa such as the kinetoplastid parasites (*e.g. Trypanosoma spp*.; Simpson 2006).

### Data filtration – Protocol_2

The widely distributed phylogenetic outlier signal in Discoba, the only major eukaryote division with no clearly identifiable stable core taxa (Fig. 3), was further explored using the sliding window method of Protocol_2 (Fig. 2). A set of 70 stable core taxa was selected based on Protocol_1 results as indicated by blue arrows along the x-axis in Figure 3. These are taxa that show minimal phylogenetic variation (low deviation scores; Fig. 3) and low levels of missing data (Fig. 4). For these analyses, the three major divisions of Discoba - Euglenozoa, Heterolobosea and Jakobida - were tested separately against the stable core taxon set.

Following a combination of automated and manual trimming, the 76 mitoP alignments vary in length from 106 to 979 positions. Using equation 2, this gives 3-10 equal length chops per protein or a total of 475 chops. Combining these chops sequentially in pairs, creates 399 consecutive sliding windows with 50% overlap, (except for terminal windows) After removing chops missing entirely for specific subgroups, the final tally is 298, 303 and 357 non-missing data chops for Jakobida, Euglenozoa and Heterolobosea, respectively.

Protocol_2 analyses of these data identify 341 potentially incongruent chops (PICs) distributed across Discoba, of which 135 are found in Jakobida, 99 in Euglenozoa and 107 in Heterolobosea (pale pink; Fig. 5). These PICs seem to be largely subgroup specific; very few are universal across Discoba (Fig. 5). All proteins identified as outliers by Protocol_1 include at least one PIC in one of the taxa for which the protein is identified as an outlier, although not necessarily all of the taxa for which the protein is an outlier (dark pink; Fig. 5). However, many more PICs are found in sequences not identified as outliers.

**Figure 5.**
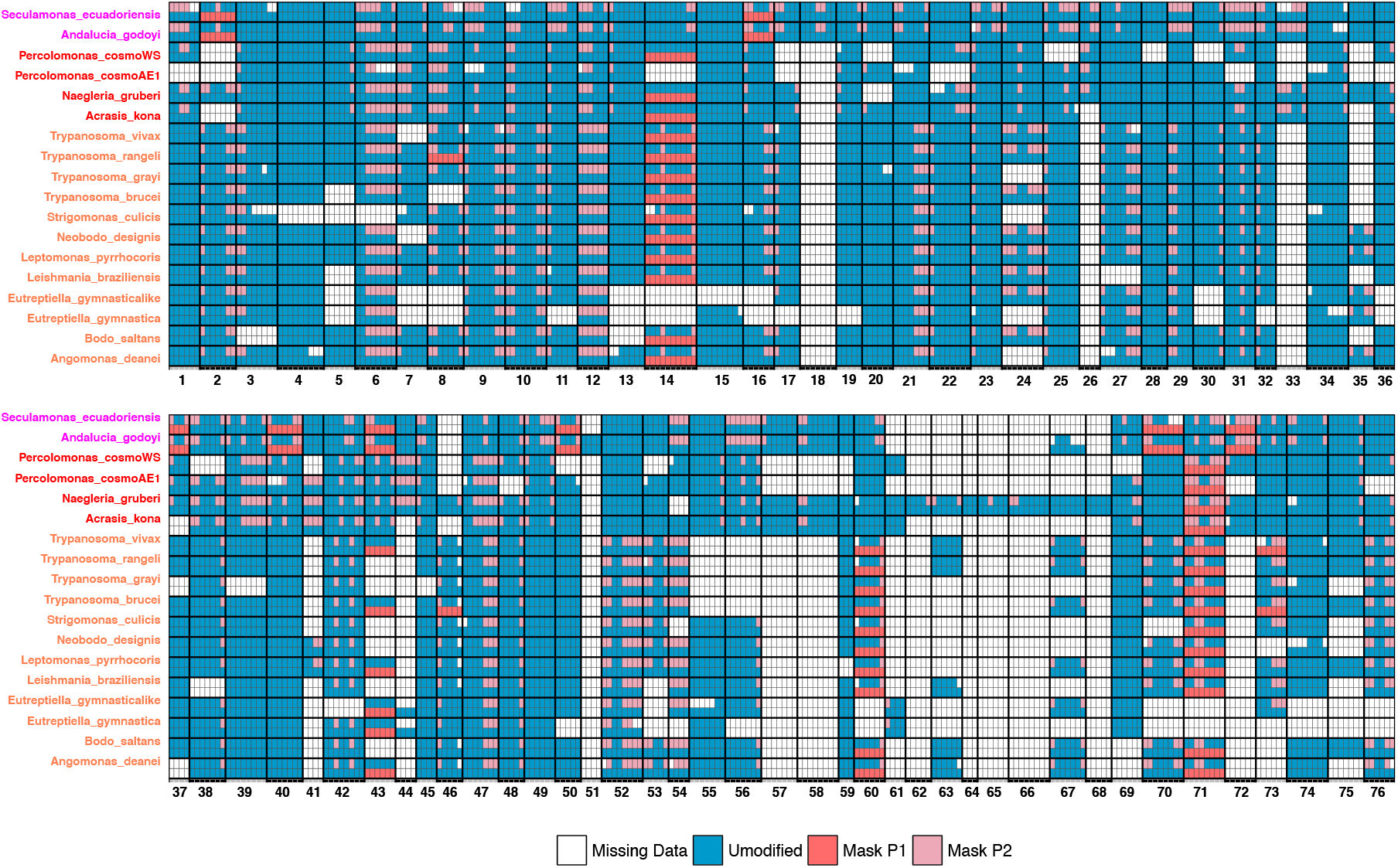
Potentially incongruent fragments of discobid mitoP proteins identified by Protocol_2. The 76 mitoP proteins were divided into 399 equal-size fragments (chops) and analyzed separately for the three major divisions of Discoba by assembling into 50% overlapping-windows followed by phylogenetic analysis (Protocol_2). Chops consistently appearing in non-canonical phylogenetic position were tagged as potentially incongruent chops (PICs). Cell colors indicate whether the chop is present (blue), absent (white), designated as an outlier by Protocol_1 (red) or as a PIC by Protocol_2 (pink). Chops are delineated by light grey lines and proteins by heavy black lines. Numbers along the x-axis indicate the protein partition number. Discobid taxon names are shown on the y-axis and color coded according to taxonomy: Jakobida (pink), Heterolobosea (red), Euglenozoa (orange).

For example, the full-length alignment for protein 40 (mtHsp70) reconstructs a monophyletic Discoba, albeit weakly and with an unresolved position among eukaryotes (Fig. S2 “full”), and a very similar topology is reconstructed by sliding windows 1 and 2 (Fig. S2 “w1-2”). In contrast, windows 3, 4, and 5 place the Heterolobosea with the Chlorophyta (green algae and land plants; Fig. S2 “w3-5”). This relationship is particularly strongly supported with the central window, window 4, which gives 88% bootstrap support for this topology (Fig. S2 “w4”). Together, these data suggest that the 2 chops that make up window 4, which also correspond to the 5’ and 3’ halves of windows 3 and 5, respectively, have a different origin from the rest of the protein in some or all of the Heterolobosea. Importantly, this anomaly is not apparent in analyses including all Discoba and the full alignment (Fig. S2 “full”), and as a result the protein does not register as an outlier for Heterolobosea under Protocol_1 (Fig. 5).

A similar phenomenon is seen with protein 03 (mtHSP60). Phylogenetic analysis of the full alignment places the Heterolobosea with the Euglenozoa, weakly grouped with Amorphea (Fig. S3a). This is essentially compatible with the unresolved position of Heterolobosea when analysed in the absence of other Discoba (Fig. S3b). However, when the Heterolobosea are represented by only chop 3 of the same protein, they appear nested within bacteria with very high support (96% bootstrap; Fig. S3c). This strongly suggests that this portion of the gene encoding mtHSP90 has an exogenous origin for some or all of the Heterolobosea.

From the Protocol_2 results, it would appear that nearly all discobid mitoP-encoding genes carry some DNA fragments of exogeneous origin. However, it is important to note that nearly half of these are “edge chops” (48.7%), meaning chops at alignment edges or bordering missing data. The relatively small number of character positions in the sliding windows makes Protocol_2 sensitive to stochastic errors, which is generally controlled for by requiring a chop to be flagged in two contiguous windows before designating it as a PIC. However, this is not possible for edge chops, which must be evaluated based only on the single window in which they occur. As a result, both type I and II errors are expected to be higher for edge chops. However, even with the exclusion of edge chops there remains a large number of potentially incongruent fragments in many discobid mitoP genes.

### Protocol_1 vs Protocol_2

Most incongruencies strong enough to be detected by both methods described here are likely to be deleted at the SGT stage. Nonetheless, the results from the two protocols show some correspondence. For example, all proteins identified as outliers by Protocol_1 include some PICs for some taxa (Fig. 5). Nonetheless, there is often only a single or a few PICs per outlier, and this is not always for the same taxon. At the same time, many PICs occur in proteins not identified as outliers by Protocol_1. These include several proteins with multiple PICs that do not appear as Protocol_1 outliers (Fig. 5). Preliminary analyses suggest that such “non-outlier multi-PIC proteins” may occur when full proteins sequences have very weak phylogenetic signal and thus escape detection in a multigene based method such as Protocol_1 (data not shown). Alternatively, such a situation could arise if a gene carried multiple PICs of different origin, such as DNA transfers from different sources, which, if fragment transfers are as common as the results here suggest (Fig. 5), would not be altogether unlikely.

Therefore, there are several reasons why it is not expected that all PICs will cause their host protein to register as an outlier. On the other hand, the fact that some outliers carry only one or a few PICs suggests that some small incongruent fragments may have a large influence. Thus, while Protocol_1 outliers are more likely to affect a multigene phylogeny, overlap between the results here suggests that both methods are effective at detecting disruptive phylogenetic signal hidden from SGTs. This is reinforced by subsequent phylogenomic analyses (below).

### Molecular phylogeny

Six versions of the mitoP data were assembled for phylogenetic analysis (Fig. 6; Table S1): the full data set of 76 mitoPs for 180 taxa (full data), the full data set masked by Protocol_1 (full, P1 mask), the 70 stable core taxa plus Discoba used as input for Protocol_2 before (core_1), the same data after Protocol_2 masking (core_1, P2 mask), and a stricter version of core_1 including only proteins with at least one sequence from each major divisions of Discoba before (core_2) and after (core_2, P2 mask2) Protocol_2 masking.

**Figure 6.**
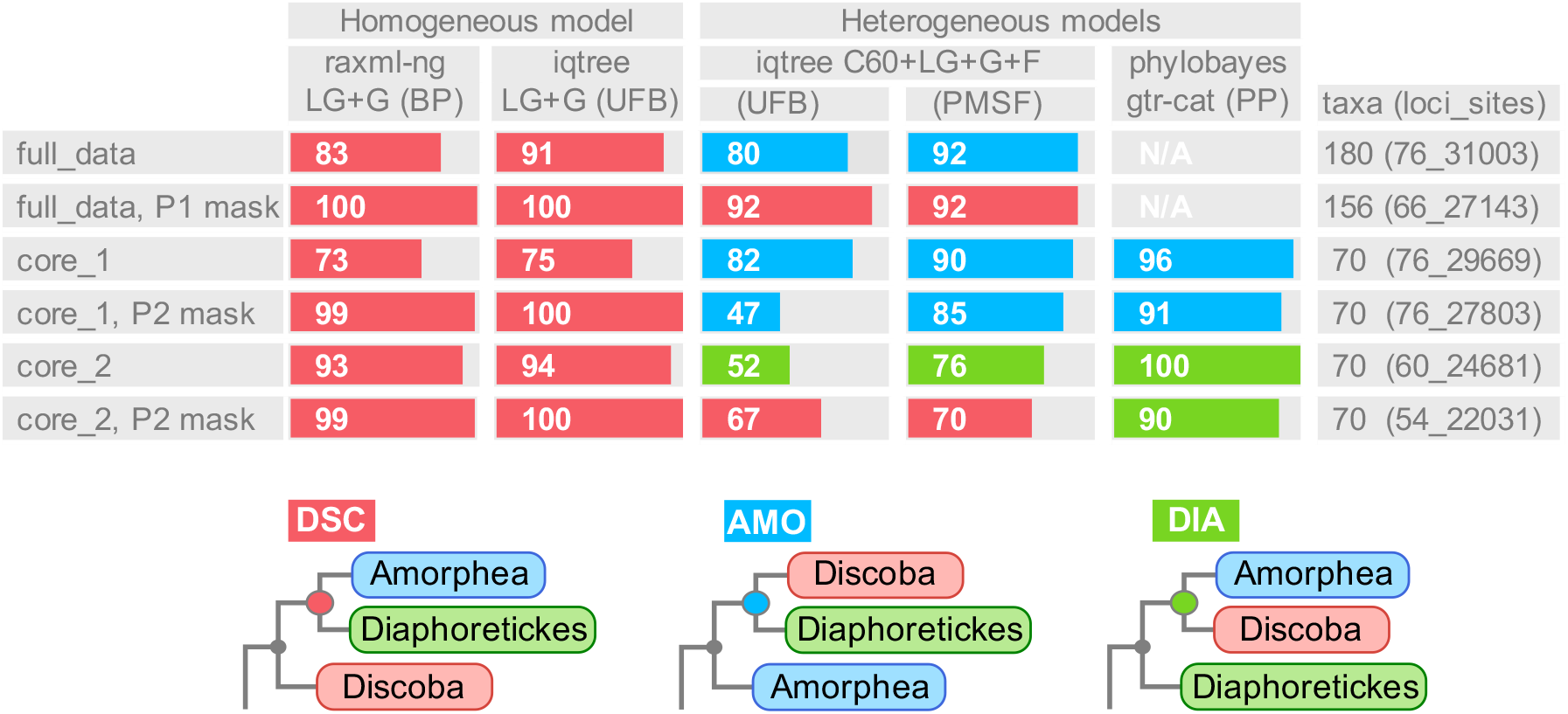
The eukaryote root based on mitoP proteins with and without masking according to Protocol_1 (persistent outliers) or Protocol_2 (incongruent fragments). Six different versions of a data set of 76 conservative mitochondrial proteins were subjected to phylogenetic analysis using five different method-model combinations to test the three main alternative eukaryote roots, shown at the bottom of the figure as cartoons and labeled with acronyms: DSC – Discoba sister (neozoan excavate root), AMO – Amorphea sister (unikont-bikont root), and DIA – Diaphoretickes sister. Support values for these roots are shown as a bar plot for six versions of the mitoP data (x-axis) analysed with five different phylogenetic method - model combinations (top y-axis). Methods are standard bootstrap with RAxML-NG and LG+4G (Raxml-NG LG-G) models (Kozlov et al. 2019), ultrafast bootstraps (UFB+bnni) using IQTREE with LG+4G (-bb) optimized with (-bnni) option, PMSF bootstraps and UFB+bnni from IQTREE using the C60 model LG+C60+F+G (Nguyen et al. 2015; Wang et al. 2018), and posterior probabilities from PhyloBayes using CAT+GTR (Laritollot et al 2012). Sizes of the mitoP data sets are shown on the right x-axis including number of taxa, number of proteins (loci) and number of aligned positions (sites). The data sets are as follows: “full_data” - the original mitoP data, “full_data, P1 mask” - full data filtered for outlier sequences using Protocol_1, “core_1” - mitoP sequences for stable core taxa plus Discoba used in Protocol_2, “core_1, P2 mask” - core data filtered for incongruent protein fragments identified by Protocol_2, “core_2” - subset of core_1 data including only proteins with at least one sequence for each subdivision of Discoba (Euglenozoa, Heterolobosea and Jakobida), and “core_2, P2 mask” - core_2 data with the same masking as core_1 P2 mask. N/A: Phylobayes analysis were conducted only for the masked and unmasked core_1 and core_2 data due to computational limitations.

These data sets were analyzed using five different combinations of maximum likelihood evolutionary models and bootstrap implementations 1) RAxML-NG with LG+gamma model and standard non-parametric bootstraps (Kozlov et al. 2019), 2) IQTREE with LG+gamma and UFB+bnni, and 3) IQTREE with the LG+C60+F+Gamma site specific (mixture) model and PSMF bootstraps and 4) UFB+bnni (Nguyen et al. 2015; Wang et al. 2018), and 5) Bayesian inference with the CAT+GTR mixture model in PhyloBayes (Lartillot et al. 2013) Due to its heavy computational demand, only the four datasets associated with Protocol_2 were analyzed by PhyloBayes.

The resulting trees are largely compatible, reproducing all well-established major divisions of eukaryotes with full support (Figs. S4-S7). The main conflicts among the trees are the position of the root, and two contentious branches within Diaphoretickes (see below). For the root, *i.e*, the branching order among the three main divisions of eukaryotes, all analyses using homogeneous or “static” models support a grouping of Amorphea and Diaphoretickes to the exclusion of Discoba. Support for this “neozoan-excavate” root reaches 99-100% bootstrap support (BP), when the data are masked according to either Protocol_1 or 2 (Fig. 6, Fig. S5-S7). In contrast, analyses of these data with mixture models find highly variable support for all possible resolutions of this trichotomy, although IQTree using the C60 model (IQTree_C60) tends to support the neozoan-excavate root with masked data, most notably data filtered according to Protocol_1 (92% BP; Fig. 6, Fig. S5).

Phylogenetic analysis using the two site heterogeneous mixture models appear to be particularly sensitive to data set composition. For example, IQTree_C60 shows strongest support for a unikont-bikont root (Amorphea sister or AMO) with the full data set (80-92% BP; Fig. 6), but analyses using Protocol_1 masked data switch completely to equally strong support for a neozoan-excavate root (Discoba sister or DIS) (92% BP; Fig. 6). PhyloBayes CAT-GTR also supports AMO with unfiltered data with a posterior probability (PP) of 0.96 (Fig. 6) but then switches to a novel Diaphoretrickes sister (DIA) with masking (core_2, P2 mask) (0.90 PP; Fig. 6, Fig. S4).

However, the shift in the PhyloBayes CAT-GTR results may not be related to the P2-masking, because the same method finds even higher support for the DIA root with the unfiltered Core_2 data (1.0 PP; Fig. 6). This suggests that it is the composition of core_2 that makes the difference for PhyloBayes CAT-GTR, and not the P2 masking. The only difference between the two datasets is 16 “discobid-poor” proteins, which are present in the core_1 data set and deleted from the core_2 data. These proteins comprise 17.0% of the total data (5051 out of 29669 alignment columns), for which Discoba are missing in 81.3% of the cells. In contrast, the core_2 data set consists only of the remaining 60 proteins for which Discoba are missing in only 15.0% of cells. The result is that the core_2 data set contains 95.7% of all the non-gap data for Discoba, while the 16 deleted proteins carry only 4.3 % of the discobid non-gap data. This, means that PhyloBayes CAT-GTR support for the AMO root is found here only with the highly discobid-depauperate data of core_1.

The other two conflicting nodes encountered here both occur within the supergroup Diaphoretickes. These are the branching order among Stramenopiles, Rhizaria and Alveolates (aka SAR) and the phylogenetic position of Cryptista (Fig. S5-S7). All analyses here tend to support a sister group relationship between Stramenopiles and Rhizaria to the exclusion of Alveolates, particularly after Protocol_1 filtering (88-94% BP; Fig. S5). The only exception to this is IQ-Tree_C60 analysis of the full unfiltered data, which places Stramenopiles and Alveolates as sister taxa (61-86% BP, data not shown). With regard to the Cryptista, all analyses with the full mitoP data reconstruct Cryptista as the sister group to red and green plants, albeit with highly variable support (46-85% BP, data not shown). However, all analyses with Protocol_1 filtered data place Cryptista as the sister to SAR (88-96% BP; Fig. S5). These are important phylogenetic questions that need to be carefully dissected in their own right, probably with more taxa in the case of Cryptista and possibly a more detailed examination with Protocol_2 in the case of SAR. It should be noted that neither node was tested with Protocol_2 filtered data as both Cryptista and Rhizaria failed the Protocol_2 stable core taxa criteria (Figs. 3 and 4).

## Discussion

Two methods (protocols) have been developed to evaluate conflict and congruence in phylogenomic data. Protocol_1 tests for persistent outliers at the whole gene/protein level using a gene jackknifing approach (Fig. 1), while Protocol_2 assesses incongruent signal at the sub-gene/protein level using a sliding-window and pre-selected core and target taxa (Fig. 2). The two methods are independent but complementary (Fig. 5). Phylogenetic analysis of 76 conservative mitochondrial proteins using the static LG-gamma model and data masked according to Protocol_1 or 2 show strong and consistent support for a neozoan-excavate root for eukaryotes (Discoba sister, 99-100% BP; Fig. 6). However, analyses of these data using two site-specific (mixture) models yield mixed results, flipping between alternative roots depending on presence/absence of masking and/or large amounts of missing data (Fig. 6). Phylogenetic analysis of Protocol_2 output also reveals strong evidence of exogenous gene fragments scattered across the mitoPs of Discoba, suggesting HGT from an assortment of donors (Fig. 5; Figs. S2 and S3).

### Two methods for congruence testing

The two methods presented here examine congruence at different levels. Protocol_1 operates at a more global scale looking for outlier signal using multigene data, while Protocol_2 operates at a more local scale using gene/protein fragments. Protocol_1 ranks sequences according to their tendency to disrupt phylogenetic signal in data subsets large enough to produce well-resolved trees but small enough to be sensitive to deviant signals. This is particularly important for detecting problems in deep phylogenetic branches, for which little signal exists in SGTs. One problem with this approach, is that many proteins simply have weak signal, e.g., if they have few variable sites, and this causes them to rank higher because their signal is easily affected by other proteins. To correct for this, each sequence-taxon data point is multiplied by the mean of the lowest 10% points (lowest deviations of the 364 samples) for that protein-taxon point. This is because in a set of subsamples where protein x is a member, the proportion of deviation contributed by x is expected to be higher in the subsamples with lower deviation values. Therefore, the final score is downweighed by the mean of the lowest 10% values, which also makes it possible for all the data points to be included in calculating the final score.

One important concern in a rooted tree is the potential presence of a substantially different evolutionary process in the outgroup. In this case bacteria are expected to have high internal phylogenetic variability of a different type and extent than that of the ingroup (eukaryotes), due to processes such as complex patterns of HGT, which is pervasive in bacteria (Juhas et al. 2009; Ke et al. 2000; Pál et al. 2005). To avoid high outgroup phylogenetic noise from overwhelming an ingroup signal, analyses are conducted separately for ingroup and outgroup. To factor in the changes in distance of the outgroup to the ingroup, the most phylogenetically consistent ingroup taxon is included in the outgroup analysis. In the case of eukaryotes this ingroup taxon is usually a plant or animal, as these include many taxa with short phylogenetic branches and low amounts of missing data.

Protocol_2 addresses the problem of incongruence at the subgene level (mosaicism). Mosaics can be difficult to detect in multigene data or even in SGTs, which suggests that most such mosaic genes, if they do exist, would probably not have a strong influence on multigene phylogeny. However, there could be cases in which mosaics are influential, for example, if multiple small HGTs were acquired from the same or closely related sources. Such a situation might occur in a cell that hosts symbionts or parasites, both of which are common in microbial eukaryotes, or even simply a microbe having a favored food source. In such cases, multiple small incongruent fragments could add up to a large influence once combined into a multigene data set. Although preliminary analyses of Discobid mitoPs did not identify such a pattern (data not shown), detecting one, if it does exist, would require more detailed analyses using smaller chops and wider taxon sampling from potential donor lineages.

Nonetheless, most persistent outliers identified by Protocol_1 are also found to include PICs by Protocol_2 (Fig. 5), and both protocols appear to be effective in masking discordant signal in phylogenomic data (Fig. 6). Thus, one potential advantage of Protocol_2 is that masking of incongruent fragments can be more precise than the usual method of masking whole genes or proteins. As a result, Protocol_2 can potentially be used to ameliorate the predicament of eliminating whole sequences if they look suspicious in SGTs or are judged to be outliers in Protocol_1, thereby leaving more data for concatenation. For example, Protocol_2 masking removes only 1866 sites or ~6% of the mitoP data, while Protocol_1 masking leads to the complete deletion of 10 mitoP proteins or (15% of the data).

Protocol_1 performs a similar function to a number of other congruence testing methods such as Conclustador and Phylo-MCOA. However, unlike Protocol_1, Conclustador (Leigh et al. 2011) does not identify which taxa are incongruent for which sequence. The software also is not complete and requires author support, which, to our knowledge is no longer available. By contrast, Phylo-MCOA (De Vienne et al. 2012) can detect taxon-partition specific incongruence. However, the method represents each partition with only a single tree and without considering the statistical significance of the tree. Therefore, the method can be misled by the stochastic errors often encountered in SGT topologies.

Of the two methods presented here, Protocol_1 is more general and easier to run. The main challenge for the user is selecting a subsample size large enough to produce meaningful trees, yet small enough that an outlier signal can still manifest. This value also depends on the amount and distribution of missing data, since each subsample must include data from all taxa. The outlier effects identified by this method are also more likely to affect the results of multigene phylogeny as they are identified on the basis of their ability to impact multigene trees. Protocol_2, on the other hand, requires informed decisions in order identify a specific set of target taxa and a phylogenetically stable but taxonomically broad core taxon set. One of the most straightforward ways to do this would be based on Protocol_1 results, as done here, but of course other sources of information could be used.

The accuracy of both Protocols is expected to increase with increased subsampling, and, as with any random sampling method, the quantity of runs is more important than optimizing their quality. Hence a fast but reasonably accurate phylogenetic method, such as fast ML with a simple model (*e.g*. RAxML with static LG) is recommended for constructing the trees, although much simpler methods like distance or unweighted parsimony do not seem to perform well (data not shown). It should be noted that both protocols are not designed to be replacements for data pre-screening with SGTs but rather aim to improve the quality of SGT pre-screened data by addressing its limitations.

### What are PICs

Protocol_2 identifies a surprisingly large number of what appear to be incongruent gene fragments in the mitoP proteins of various Discoba (Fig 5), some examples of which are confirmed by molecular phylogeny (e.g., Figs. S2, S3). While many discobid PICs may be false positives, particularly edge PICs, these are still only roughly half of the PICs identified here. Moreover, it is unlikely that all or even most edge PICs are false. Thus, it appears that a substantial number of discobid mitoP-encoding genes are mosaic. Such mosaicism could result from recombination of host genes with paralogs elsewhere in their genomes or with externally derived gene fragments (xenologs). Studies of HGT tend to focus on whole genes, whose acquisition could allow a new host to acquire a novel function, a process that appears to be pervasive in bacteria (Hall et al. 2017). However, there is also the potential that, if the host already has a homolog of the “invading” DNA, the host and exogenous sequences could recombine and create a mosaic gene.

Theoretically, there should be considerable potential for random partial gene replacement for the bacterial-like genes of microbial eukaryotes, as many eukaryotic microbes are bacteria predators and/or host bacterial symbionts or parasites. Any of these could potentially break down and release random genetic material into the host cell (Husnik and McCutcheon 2018). Moreover, the largest class of bacterial-like genes in eukaryotes are genes encoding proteins targeted to organelles of endosymbiotic origin (Ku et al. 2015). Such mosaic gene have been found in plant mitochondrial DNA, albeit from other plants and potentially mediated by viruses (Richardson & Palmer 2007). The results presented here suggest that partial gene replacement is more pervasive in eukaryotes, at least for some taxa and some sets of genes, in this case, nuclear genes encoding bacterial-like mitochondrial proteins of Discoba (Fig. 5).

### Deep phylogeny of eukaryotes based on mitoP data

The mitoP data used here have been the subject of three previous studies, abbreviated here as DL2012, D2015 and H2014 (Derelle et al. 2015; Derelle & Lang 2012; He et al. 2014). Despite using partly overlapping data these studies found very different results with DL2012 and D2015 supporting the unikont-bikont or Amorphea sister root and H2014 the neozoan-excavate or Discoba sister root. In the current work, all proteins used in the previous analyses were combined and treated equally to build a new dataset, also adding several more recently sequenced taxa. These data were then objectively and systematically screened for persistent outliers (Protocol_1; Fig. 1) and incongruent fragments (Protocol_2: Fig. 2) using no assumption on the position of the root. Trees were then constructed from these data before and after protocol-based masking and before and after removing the most discobid-depleted proteins (Fig. 6).

The single consistent result obtained here for the eukaryote root is a neozoan-excavate root, which specifies Amophea+Diaphoreticks (“neozoa”) and Discoba (“excavata”) as two separate major divisions of eukaryotes. This result is fully consistent using the LG+GAMMA model, which finds moderate support for this root with the full unfiltered data set (73-94% BP; Fig. 6) and full support after either Protocol_1 or Protocol_2 filtering (99-100% BP; Fig. 6). In contrast, very inconsistent results were obtained with two mixture model-based methods. Together, these switch among all three possible solutions to what is essentially an unresolved trichotomy depending on presence/absence of filtering and/or large amounts of discobid missing data (Fig. 6). This suggests there is a problem with applying mixture models to these data. This seems particularly true for PhyloBayes CAT-GTR, also the only method-model combination strongly supporting an AMO root in previous studies (DL2012, D2015).

### The cost of complexity

Model selection has become something of a holy grail in phylogenetics, although recent studies indicate that better fitting models (as judged by criteria such as AIC or BIC) do not necessary produce more reliable trees with either DNA (Abadi et al 2019) or protein sequences (Spielman 2020). Model complexity and error are direct trade-offs, because each model parameter value is an estimate with an associated error (Huelsenbeck & Crandall 1997). Thus, increasingly complex models run the risk of over-fitting, whereby the model fits increasingly well to a smaller subset of the data. An added complication of highly complex models such as C60 and especially CAT-GTR is that they are computationally heavily demanding. Therefore, these models are still largely unexplored in terms of their performance with various problematic data types such as persistent outliers, varying quantities and distributions of missing data (*e.g*., patterned missing data) or model variation across the tree (heterotachy). In fact, recent work has indicated that an access of categories in mixture models can be problematic (Li et al. bioRxiv).

In the case of the mitoP data, the inconsistency of results using mixture models is particularly apparent when contrasted between before and after protocol-based masking. As expected, such masking increases nodal support in analyses using simpler models (Fig. 6). However, it causes radical shifts in topology and support with both mixture models. For example, IQTree_C60 finds 92% support for an AMO root with unfiltered data, but then switches to 92% support for a DSC root after Protocol_1 filtering. Moreover, with core data (most stable taxa plus Discoba), IQTree C60 finds all three possible roots depending on masking and core group composition. PhyloBayes CAT-GTR, on the other hand may be especially sensitive to missing data as it supports an AMO root with core_1 taxa, with or without masking, but a DIA root with reduced missing data, again with or without masking (Fig. 6).

### Missing taxa

Many taxa are not included in these analyses presented here, including newly discovered species that may represent novel major divisions of eukaryotes (Burki et al. 2020). However, none of these taxa are widely sampled or have sufficient genomic data to allow reliable pre-screening for phylogenomics. We chose to exclude these taxa to focus on a single specific question, the branching order among Amorphea, Diaphoretickes and Discoba. Thus, we cannot rule out the possibility that one or more of the excluded lineages represents a closer sister lineage to Amorphea+Diaphoretickes or Discoba or a deeper branching sister group to all three. We also obviously cannot rule out the possibility that one or more such organisms or taxon groups will be discovered in the future, and hopefully they will be. However, no addition of taxa should affect the fundamental relationship described here, that Amorphea and Diaphoretickes are more closely related to each other than either is to Discoba. Careful selection of data and taxa is of paramount importance to accurate phylogenetic reconstruction and removing strong outlier data is crucial to improve tree accuracy (Philippe et al. 2017), particularly for the deepest branches. Moreover, precisely defining relationships among known major taxa will make it easier to assign novel taxa to their correct position in the tree.

## Conclusions

Two new protocols are presented for identifying incongruence in multigene data at the level of whole (Protocol_1) and subgene (Protocol_2) levels. Protocol_1 is quite easy to apply using scripts archived in GitHub (https://github.com/caeu/eukRoot). Protocol_2 requires considerably more user expertise including clearly defined hypotheses. However, it can potentially detect mosaic genes, and the results reported here suggest that these are more widespread in eukaryotes than previously known (Bergthorsson et al. 2003; Bock 2010; Mower et al. 2004; Nielsen et al. 1998; Richardson & Palmer 2007). Most importantly, application of these protocols to a data set of bacterial-like mitochondrial proteins and subsequent phylogenetic analyses place the Discoba as a sister clade to Amorphea plus Diaphoretickes, with no consistent support found with these data for any alternative hypothesis. Probably the most important missing taxa here are the remaining “excavates”, the metamonads or “de-mitochondriate excavates”, which, due to their lack of respiring mitochondria also lack nearly all core mitoPs. Thus, the neozoan-excavate root is just one step, albeit an important one, in defining the root of the eukaryote tree.

## Acknowledgement

This work was supported by VR grant (Project VR 2017-04351) and Project SNIC2019/3-266 from the Uppsala Multidisciplinary Center for Advanced Computational Science (UPPMAX). The authors thank Lars Arvestad and Mikael Thollesson for helpful comments on the manuscript.

